# Vitrification after multiple rounds of sample application and blotting improves particle density on cryo-electron microscopy grids

**DOI:** 10.1101/094623

**Authors:** Joost Snijder, Andrew J. Borst, Annie Dosey, Alexandra C. Walls, Anika Burrell, Vijay S. Reddy, Justin M. Kollman, David Veesler

**Affiliations:** Department of Biochemistry, University of Washington, Seattle, Washington, USA.; Department of Integrative Computational and Structural Biology, The Scripps research Institute, La Jolla, California, USA.

## Abstract

Single particle cryo-electron microscopy (cryoEM) is becoming widely adopted as a tool for structural characterization of biomolecules at near-atomic resolution. Vitrification of the sample to obtain a dense distribution of particles within a single field of view remains a major bottleneck for the success of such experiments. Here, we describe a simple and cost-effective method to increase the density of frozen-hydrated particles on grids with holey carbon support films. It relies on performing multiple rounds of sample application and blotting prior to plunge freezing in liquid ethane. We show that this approach is generally applicable and significantly increases particle density for a range of samples, such as small protein complexes, viruses and filamentous assemblies. The method is versatile, easy to implement, minimizes sample requirements and can enable characterization of samples that would otherwise resist structural studies using single particle cryoEM.

---

Single particle cryo-electron microscopy (cryoEM) is becoming an increasingly important tool in structural biology (Nogales and Scheres 2015; Cheng *et al.* 2015). Due to recent advances in detector efficiency, microscope stability and image processing, the 3D structure of biomolecular complexes spanning a wide range of sizes, from haemoglobin to microtubules and intact viruses, can be reconstructed at (near-) atomic resolution (Bai *et al.* 2013; Brilot *et al.* 2012; Campbell *et al.* 2012; Campbell *et al.* 2015; Koshouei *et al.* 2016; Liu *et al.* 2010; Nogales 2015; Scheres 2012; Walls *et al.* 2016a; Walls *et al.* 2016b).

Whereas sample requirements are less restrictive in cryoEM compared to X-ray crystallography or nuclear magnetic resonance, successful vitrification of protein complexes remains a major bottleneck (Grassucci *et al.* 2007). A solution of the particles under study is pipetted onto holey grids made of carbon-coated copper or gold (Quispe *et al* 2007; Russo *et al* 2014). The holes in the grid support film become filled with particles suspended in buffer and the excess solution is blotted away using filter paper. The grid is then immediately plunged into a bath of liquid ethane, maintained at liquid nitrogen temperature (−180°C) to vitrify the protein solution and preserve it in a near-native frozen-hydrated state (Dubochet 1988). Numerous specimens, however, are reluctant to populate the holes and remain adsorbed on the carbon support film instead, even after significantly increasing the sample concentration. Furthermore, many protein complexes are precious, due to the difficulty to purify them in large quantities, such that using higher concentrations is not an option.

In an effort to improve the applicability of single particle cryoEM, many strategies have been developed for sample preparation. For instance, samples can be deposited on 2D crystals of streptavidin, captured on immuno-affinity supports or on a continuous carbon or graphene oxide film, prior to plunge-freezing in liquid ethane (Han *et al.* 2012; Kelly *et al.* 2008; Pantelic *et al.* 2010; Wagenknegt *et al.* 1988; Yu *et al.* 2016). However, those methods can be costly, technically challenging, reduce the contrast of the images (except for graphene oxide) and may introduce structural defects related to adsorption on a surface.

Here, we present a simple and cost-effective method that builds on the standard plunge freezing procedure to overcome the aforementioned shortcomings. Although previous reports mentioned the possibility of multiple sample applications to increase particle density in suspended vitreous ice (Cheng *et al.* 2015), this has never been described in detail to the best of our knowledge. We systematically tested this method on various samples and consistently observed that it greatly improved particle density in thin films of suspended vitreous ice, including cases in which ramping up the concentration failed to do so. We believe this procedure can benefit the exponentially growing numbers of single particle cryoEM practitioners who will inevitably run into samples that prove reluctant to evenly spread over holey cryoEM grids.

This method involves performing several successive rounds of sample application and blotting before plunge freezing. We tested up to four blotting rounds and observed a marked additive effect of particle density at each step. All sample applications and blotting steps can be carried out on the lab bench, similarly to what is done for negative staining sample preparation, until the final sample application, at which point the tweezers are mounted in a plunge-freezing apparatus to perform the final blotting step using standard procedures. Alternatively, all rounds of sample application and blotting can be performed with the tweezers mounted in a plunge-freezing apparatus, if desired. We used Whatman No. 1 filter paper, which is the only extra requirement over our standard grid preparation procedure. For all examples that follow, samples were vitrified using Protochips C-flat grids and imaged on an FEI T12 electron microscope equipped with a Gatan US4000 CCD camera, an FEI TF20 or an FEI Titan Krios, equipped with a Gatan K2 direct detector (Li *et al.* 2013). We used MotionCorr2 (Zheng *et al.* 2016) for movie frame alignment and binned all images by a factor of 4 for visualization.

To demonstrate the efficacy of this method, we used a computationally designed octahedral protein cage (King *et al.* 2012), assembled from 24 identical subunits, as a test specimen (O3-33, Figure 1). At 1 mg/mL, the O3-33 solution yielded an optimal particle density for single particle cryoEM imaging. Diluting the sample five-or tenfold significantly reduced the distribution of particles over suspended ice. This effect could be partially overcome for both dilutions after a second round of sample application and blotting. After the fourth blotting step, the particle density in suspended vitreous ice became comparable to that observed with grids prepared with a single application of a ten times higher protein concentration. These results illustrate that the method can greatly alleviate sample requirements for single particle cryoEM, both in terms of protein concentration and total amount required.

**Figure 1.**
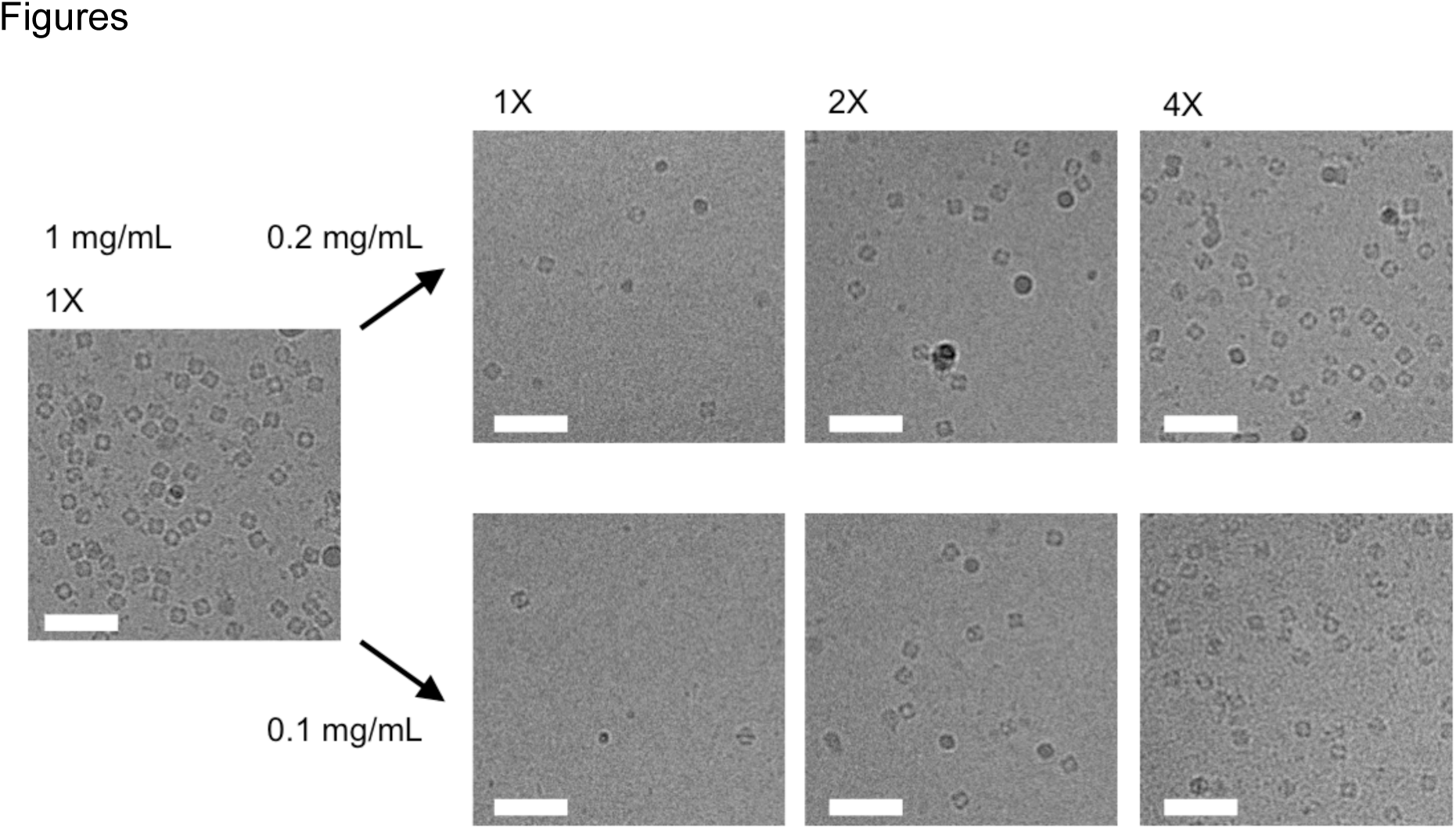
The effect of multiple rounds of sample application and blotting on particle density in thin films of suspended vitreous ice observed with samples of O3-33. Sample concentrations and number of application steps are indicated. Scale bars: 50 nm.

To assess the versatility of this approach, we implemented this multiple blotting scheme for a range of protein complexes, several of which had proven reluctant to the standard vitrification procedure.

Adenoviruses are ∼1000 Å-wide non-enveloped icosahedral pathogens involved in respiratory, ocular and gastro-intestinal infections (Veesler *et al* 2014). Samples of adenovirus serotype 26 virions showed a low propensity to evenly spread in thin films of suspended vitreous ice, which was problematic because a limited amount and concentration of sample was available for our cryoEM study. Using a double blotting strategy, we were able to overcome this issue, as attested by the densely packed virions in the micrograph presented in Figure 2, and to collect a large dataset which led to the determination of a structure at 3.7 Å resolution (manuscript in preparation).

**Figure 2.**
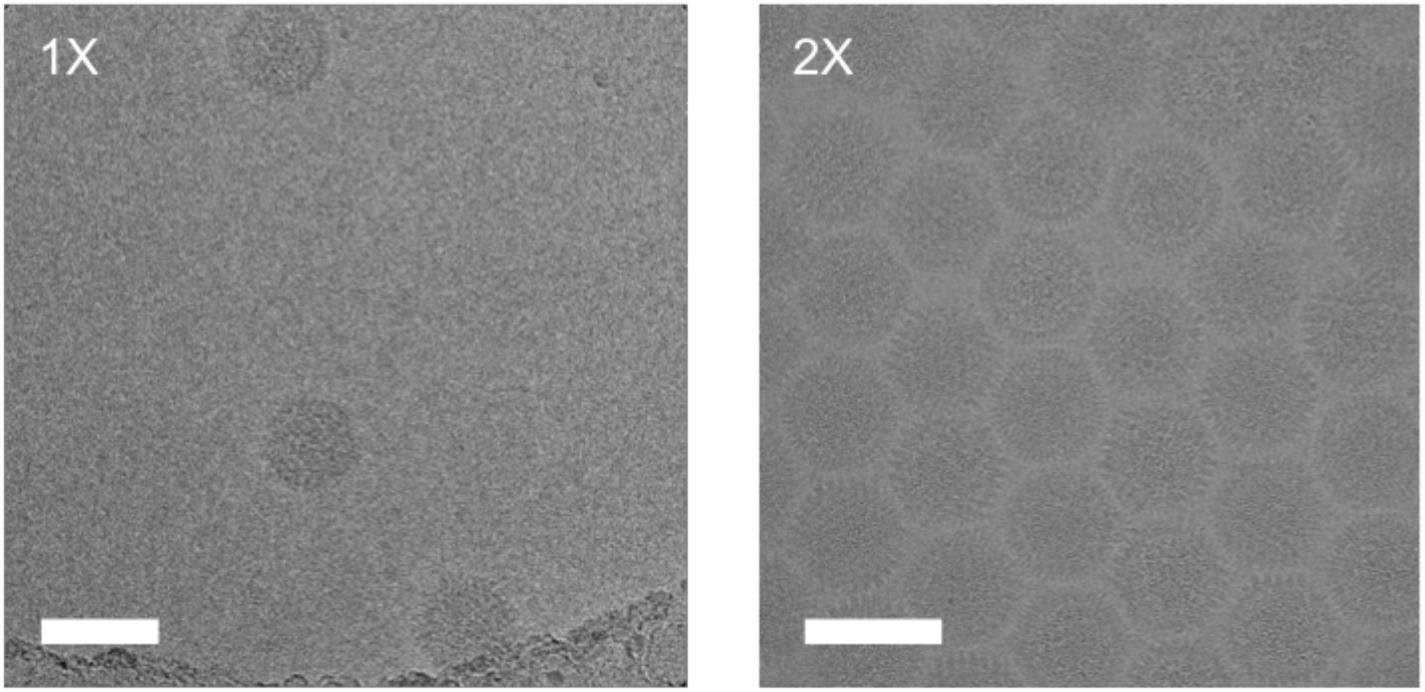
Multiple rounds of sample application and blotting of human adenovirus. Scale bars: 100 nm.

Yeast glucokinase 1 and human inosine-5’-monophosphate dehydrogenase are two key metabolic enzymes involved in carbohydrate and nucleotide metabolism, respectively. These two enzymes self-assemble as elongated homomeric filaments, which can only be structurally characterized by cryoEM due to their repetitive and flexible nature. Although sample availability was not a limiting factor for these two specimens, both proved reluctant to spread over the holes of cryoEM grids. We successfully enhanced particle distribution, following up to three consecutive applications of sample onto the grids, to achieve a particle density that was useful for high-resolution imaging (Figures 3 and 4).

**Figure 3.**
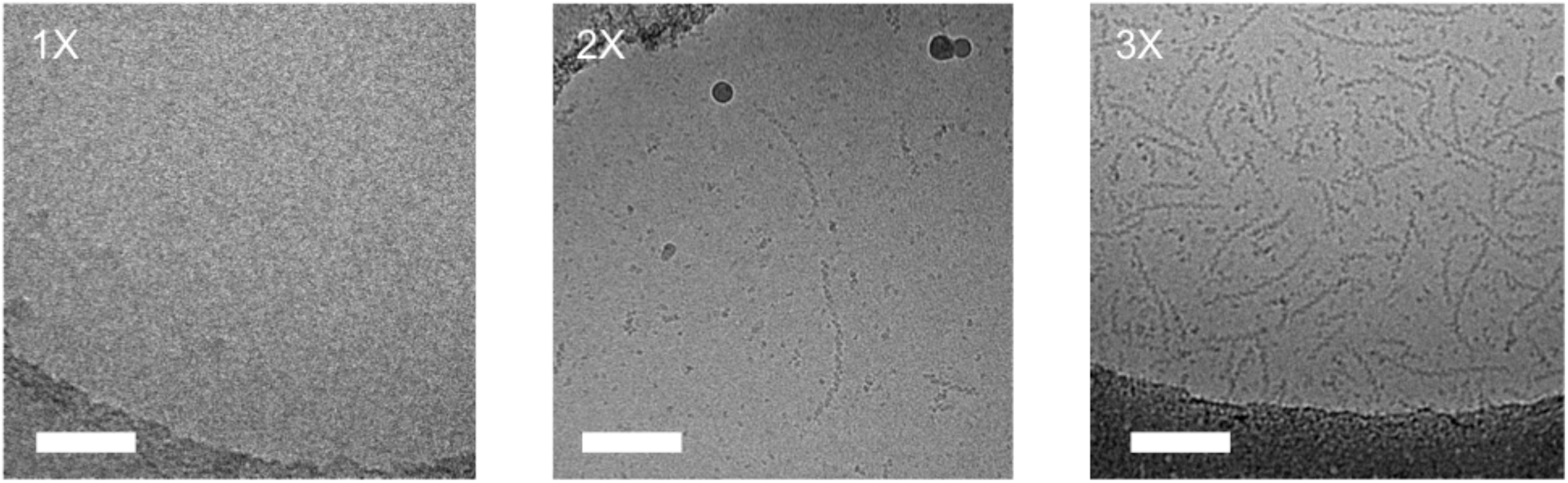
Multiple rounds of sample application and blotting of filaments of yeast glucokinase-1 in the presence of ATP and glucose. Scale bars: 100 nm.

**Figure 4.**
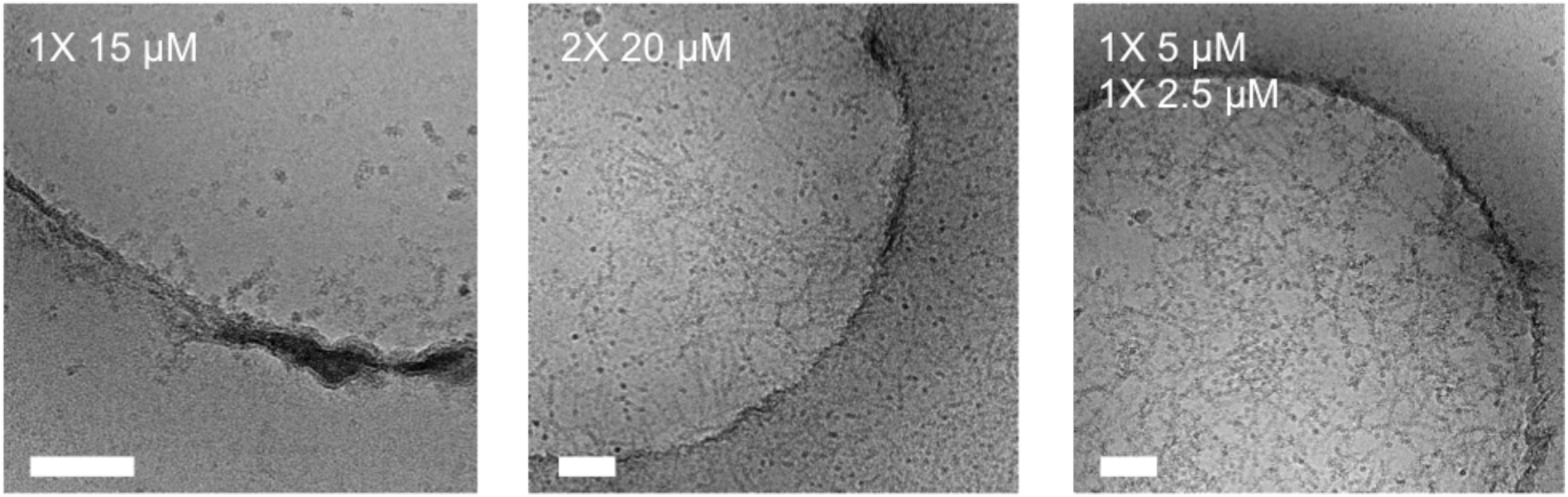
Multiple rounds of sample application and blotting of filaments of inosine-5’-monophosphate dehydrogenase. Scale bars: 100 nm.

Detergents or amphipols are oftentimes used at low concentrations to supplement a protein solution prior to grid preparation with the aim of reducing aggregation and/or promoting more isotropic particle orientations (Lyumkis *et al.* 2013; Chowdury *et al.* 2015). However, this strategy requires using much higher concentrations than one would need in the absence of such additives to ensure the sample will uniformly spread over holey carbon grids. Similar problems are encountered in cryoEM studies of detergent or amphipol solubilized membrane proteins.

We used the mouse hepatitis virus (MHV) spike (S) glycoprotein ectodomain (Walls *et al.* 2016a; Walls *et al* 2016c) to illustrate the usefulness of our approach when vitrifying samples in presence of detergents (see Figure 5). A single round of sample application and blotting of the MHV S ectodomain in the presence of 0.01% NP40 yielded few particles in suspended vitreous ice, even at concentrations up to 4 mg/mL. After the second round of blotting, many more particles could be observed in the ice. We would like to emphasize that in the absence of detergent, a single sample application resulted in a very dense distribution of particles, even at less than half the concentration. We obtained similar results with the HIV envelope glycoprotein ectodomain in the presence of 85 μM of dodecyl-β-D-maltoside (a sample known to be prone to aggregation upon vitrification, Lyumkis *et al* 2013). Although few particles were detected in vitreous ice after a single round of blotting, good particle density was achieved after the second sample application, thereby alleviating the need for large amount of protein (Figure 6).

**Figure 5.**
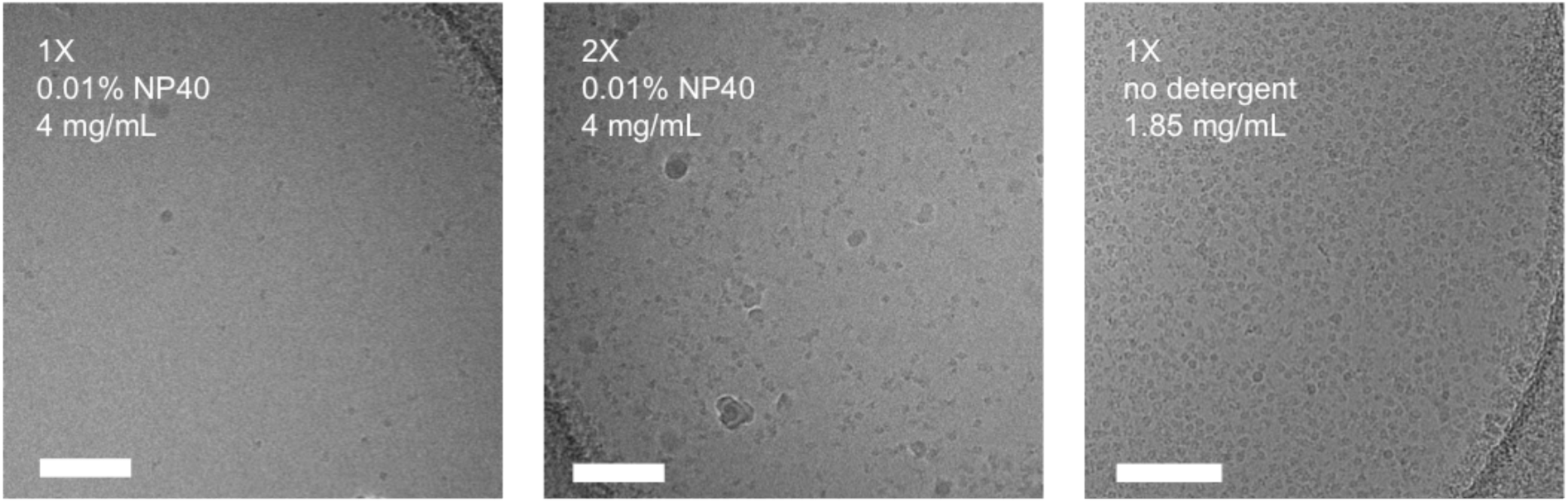
Multiple rounds of sample application and blotting of coronavirus MHV spike glycoprotein ectodomain in the presence of 0.01% NP40. Scale bars: 100 nm.

**Figure 6.**
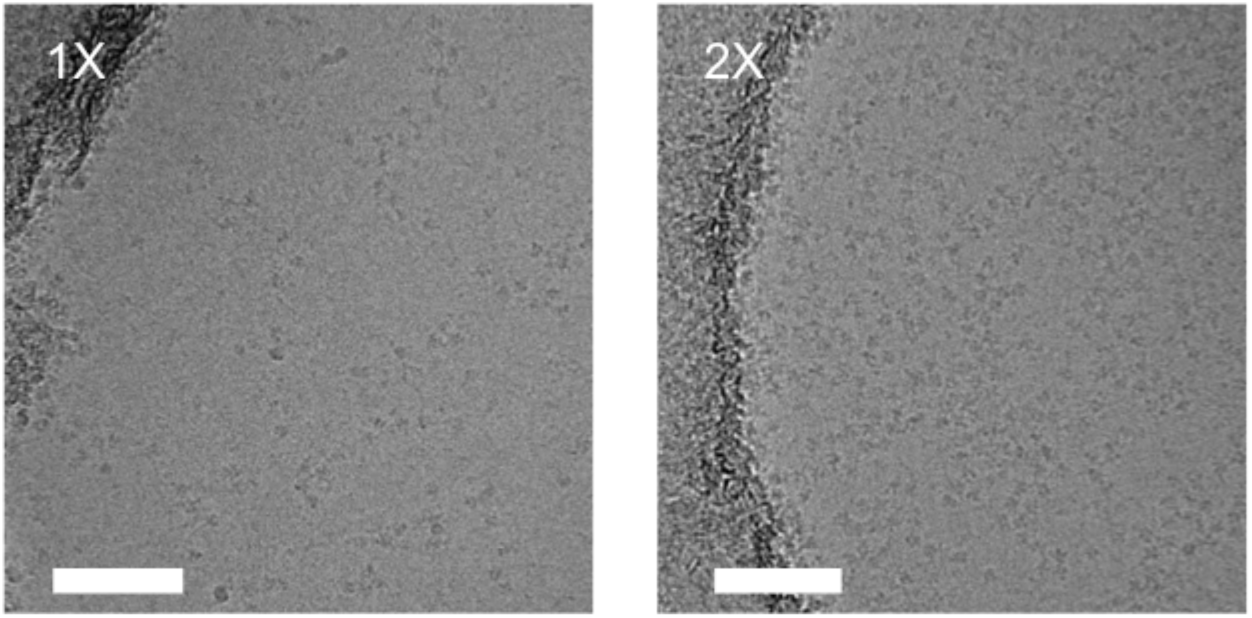
Multiple rounds of sample application and blotting of HIV envelope glycoprotein ectodomain in the presence of 85 μM dodecyl-maltoside. Scale bars: 100 nm.

In conclusion, using a simple and cost-effective trick, one can increase the applicability of single particle cryoEM to protein complexes that cannot be purified in large quantities and/or that behave poorly upon vitrification. Multiple steps of sample application can force particles into the vitreous iced-filled holes of a cryoEM grid to minimize sample requirements for structural studies.

## Acknowledgements

Research reported in this publication was supported by the National Institute of General Medical Sciences (NiGMS) under award number 1R01GM120553-01 (D.V.), 1R01GM118396-01 (to J.M.K.), R01AI070771 to VSR, T32GM008268 (to A.C.W.) and T32GM008268 (to A.J.B.). J.S. acknowledges support from the Netherlands Organization for Scientific Research (NWO, Rubicon 019.2015.2.310.006) and the European Molecular Biology Organisation (EMBO, ALTF 933-2015). We are grateful to Neil King (University of Washington) for providing the O3-33 construct.

